# Anti-proliferative therapy for HIV cure: a compound interest approach

**DOI:** 10.1101/063305

**Authors:** Daniel B. Reeves, Elizabeth R. Duke, Sean M. Hughes, Martin Prlic, Florian Hladik, Joshua T. Schiffer

## Abstract

In the era of antiretroviral therapy (ART), HIV-1 infection is no longer tantamount to early death. Yet the benefits of treatment are available only to those who can access, afford, and tolerate taking daily pills. True cure is challenged by HIV latency, the ability of chromosomally integrated virus to persist within memory CD4^+^ T cells in a non-replicative state and activate when ART is discontinued. Using a mathematical model of HIV dynamics, we demonstrate that treatment strategies offering modest but continual enhancement of reservoir clearance rates result in faster cure than abrupt, one-time reductions in reservoir size. We frame this concept in terms of compounding interest: small changes in interest rate drastically improve returns over time. On ART, latent cell proliferation rates are orders of magnitude larger than activation and new infection rates. Contingent on subtypes of cells that may make up the reservoir and their respective proliferation rates, our model predicts that coupling clinically available, anti-proliferative therapies with ART could result in functional cure within 2-10 years rather than several decades on ART alone.

## Introduction

The most significant accomplishment in HIV medicine is the suppression of viral replication and prevention of AIDS with antiretroviral therapy (ART). However, HIV cure remains elusive due to viral latency, the ability of integrated virus to persist for decades within CD4^+^ T cells in a latent state. When ART is discontinued, latent cells soon activate, and virus rebounds [1, 2]. HIV cure strategies aim to eradicate the latent reservoir of infected cells [3] but have been unsuccessful except in one notable example [4]. In addition, substantial technological and financial hurdles preclude the widespread use of many developing cure strategies. The anti-proliferative therapies we propose here are used widely, permitting broad and immediate availability following a proof of efficacy study.

Several recent studies link cellular proliferation (both antigen-driven expansion and homeostatic proliferation) with persistence of the HIV reservoir on long-term ART (> 1 year) [5, 6, 7, 8, 9, 10, 11, 12, 13]. Using a mathematical model, we demonstrate that continuous, modest reductions in latent cell proliferation rates would deplete the latent reservoir more rapidly than comparable increases in HIV activation as occurs with latency reversing agents. Further, we find that more rapid reservoir elimination on anti-proliferative therapy occurs with lower pre-treatment reservoir size and higher proportions of rapidly proliferating effector and central memory CD4^+^ T cells in the reservoir.

Based on analogies to finance, we call this strategy “compound interest cure.” We demonstrate the promise of the compound interest approach by identifying reservoir reduction commensurate with predictions from our model in HIV-infected patients treated with mycophenolate mofetil (MMF) in past studies. We confirm the anti-proliferative effect of MMF on naoïve and memory CD4^+^ T cell subsets via *in vitro* experiments.

## Results

### ART decouples latent pool dynamics from ongoing infection

Our model is visualized in Fig. 1 and detailed in the Methods. If ART is perfectly effective, all susceptible cells are protected from new infection, even when cells activate from latency. Thus, the dynamics of the latent cells can be considered separately, decoupled from the dynamics of the other cell types, and the only mechanisms changing the latent cell pool size are cell proliferation, death, and activation (bottom panel, Fig. 1).

**Figure 1:**
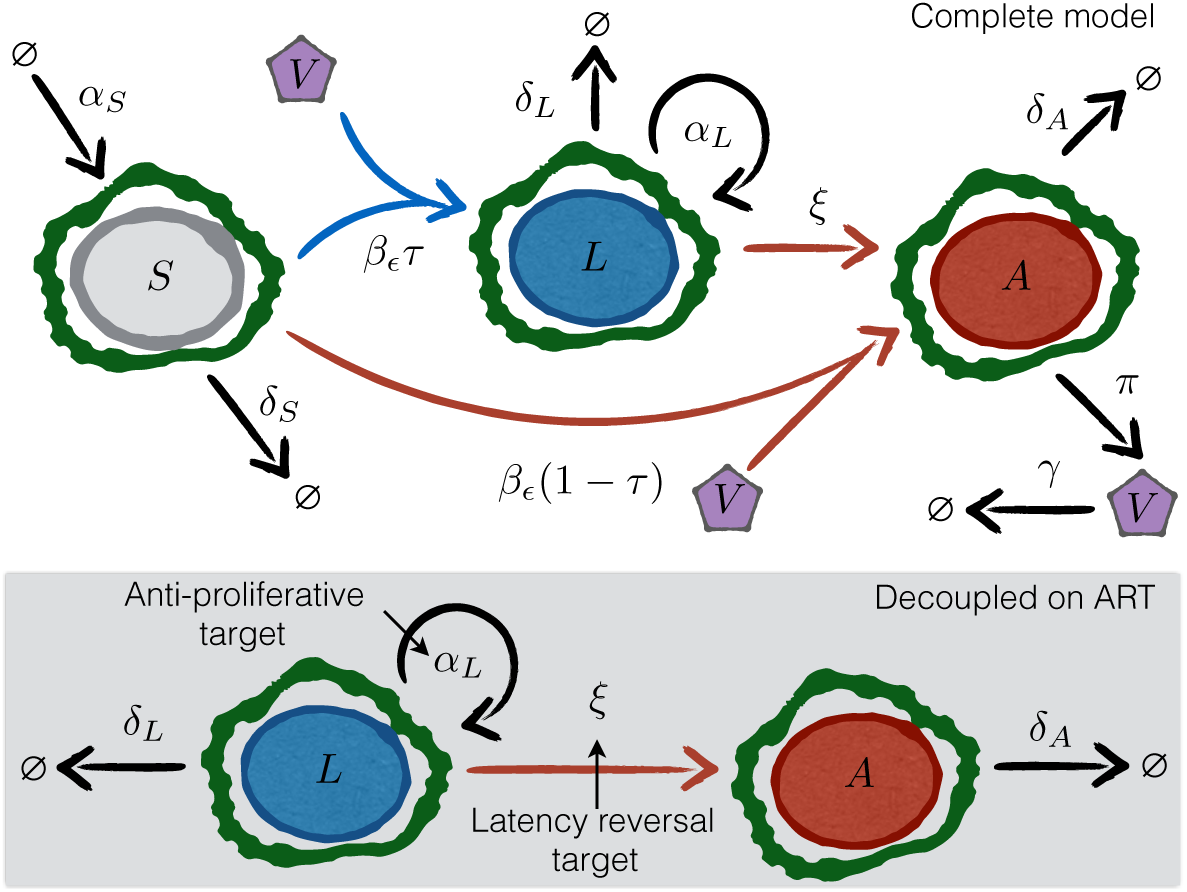
Schematics of models for HIV dynamics on and off ART. The top panel shows all possible transitions in the model (equation (1)). The bottom shaded panel shows the available transitions for the decoupled dynamic equations when ART suppresses the virus. Model parameters are given in Table 1. HIV virus *V* infects susceptible cells *S* at rate *β* reduced by ART of efficacy *ϵ* to *β*_*ϵ*_. The probability of latency given infection is *τ*. The rate of activation from latently infected cells (*L*) to actively infected cells (*A*) is *ξ*. Cellular proliferation and death are determined by rates *α* and *δ* for each compartment. The mechanisms of action of anti-proliferative and latency reversal therapies are to decrease *α*_*L*_ and increase *ξ*, respectively.

However, perfectly effective ART is not strictly necessary to consider the latent pool separately. As previously described[14, 15], we define ART “critical efficacy” *ϵ*_*c*_ as the ART efficacy above which there is no set-point viral load, *i.e.* virus decreases rapidly with time (see Methods). Above the critical efficacy, viral production from activation could cause some new cell infection, but because the probability of latency (*τ*) is so low, new infection does not affect reservoir size or dynamics meaningfully. Using parameters from Table 1, we find *ϵ*_*c*_ *∼* 85%. Because true ART efficacy is generally greater than this efficacy [16], we predict little *de novo* infection in ART-suppressed patients, consistent with the lack of viral evolution following years of ART without re-seeding of the latent reservoir[17, 10, 11, 8, 13].

**Table 1:** Parameters used in the HIV latency model. All cellular rates are for CD4^+^ T cells. *Death rates for each cell type are calculated using the total clearance as *δ*_*i*_ = *α*_*i*_ − *ξ* − *θ*_*L*_ with *i* ∈ [*L*, *cm*, *em*, *n*]

| Parameter | Value | Dimensions | Source | Meaning |
| --- | --- | --- | --- | --- |
| $\theta_L$ | $-5.2 \times 10^{-4}$ | $\text{day}^{-1}$ | [1, 26] | net latent clearance rate on ART |
| $\delta_L$ | 0.0155 | $\text{day}^{-1}$ | calculated* | latent central memory cell death rate |
| $\alpha_L$ | 0.015 | $\text{day}^{-1}$ | [54] | latent cell proliferation rate |
| $\alpha_{cm}$ | 0.015 | $\text{day}^{-1}$ | [54] | latent central memory cell $T_{cm}$ proliferation rate |
| $\alpha_{em}$ | 0.047 | $\text{day}^{-1}$ | [54] | latent effector memory cell $T_{em}$ proliferation rate |
| $\alpha_n$ | 0.002 | $\text{day}^{-1}$ | [54] | latent naïve cell $T_n$ proliferation rate |
| $\xi$ | $5.7 \times 10^{-5}$ | $\text{day}^{-1}$ | [21] | activation rate |
| $\alpha_A$ | 0 | $\text{day}^{-1}$ | [45] | active proliferation rate |
| $\delta_A$ | 1.0 | $\text{day}^{-1}$ | [61] | active death rate |
| $\tau$ | $10^{-4}$ | $\text{day}^{-1}$ | [29] | probability of latency given infection |
| $\alpha_S$ | 300 | $\text{cells}/(\mu\text{L-day})$ | [56] | susceptible growth rate |
| $\delta_S$ | 0.2 | $\text{day}^{-1}$ | [56] | susceptible death rate |
| $\beta$ | $10^{-4}$ | $\mu\text{L}/(\text{virus-day})$ | [56, 45] | HIV infectivity off ART |
| $\beta_\epsilon$ | $\beta(1 - \epsilon)$ | $\mu\text{L}/(\text{virus-day})$ | [14, 15] | HIV infectivity on ART (with efficacy $\epsilon \in [0, 1]$) |
| $\pi$ | $10^3$ | $\text{virus}/(\text{cell-day})$ | [63, 45] | viral production rate |
| $\gamma$ | 23 | $\text{day}^{-1}$ | [60] | viral clearance rate |

### Sustained mild effects on clearance rate deplete the reservoir more rapidly than large, one-time reservoir reductions

The HIV cure strategy most extensively tested in humans is “shock-and-kill” therapy: latency reversing agents activate HIV in latent cells to replicate and express HIV proteins, allowing immune clearance while ART prevents further infection [3]. Other strategies in development include therapeutic vaccines [18], viral delivery of DNA cleavage enzymes [19], and transplantation of modified HIV-resistant cells [20] informed by the “Berlin patient” [4]. Some of these therapies manifest as one-time reductions in the number of latent cells. We simulate such instantaneous decreases using equation (4) and cure thresholds described in Methods. Briefly, using ART interruption data, Hill *et al.* and Pinkevych *et al.* estimated the number of latently infected cells that would result in ART-free suppression of viremia for one year (Hill 1-yr and Pinkevych 1-yr) versus 30 years, Hill cure (Hc), in 50% of HIV-infected patients [21, 22]. With the reservoir clearance rate *θ*_*L*_ constant and a 100-fold reduction in reservoir size *L*_0_, the Pinkevych 1-yr threshold is immediately satisfied, but the Hill 1-yr and Pinkevych cure still require 15 years of ART. Hill cure requires a 1,000-fold reduction and more than 10 subsequent years of ART (Fig. 2a).

**Figure 2:**
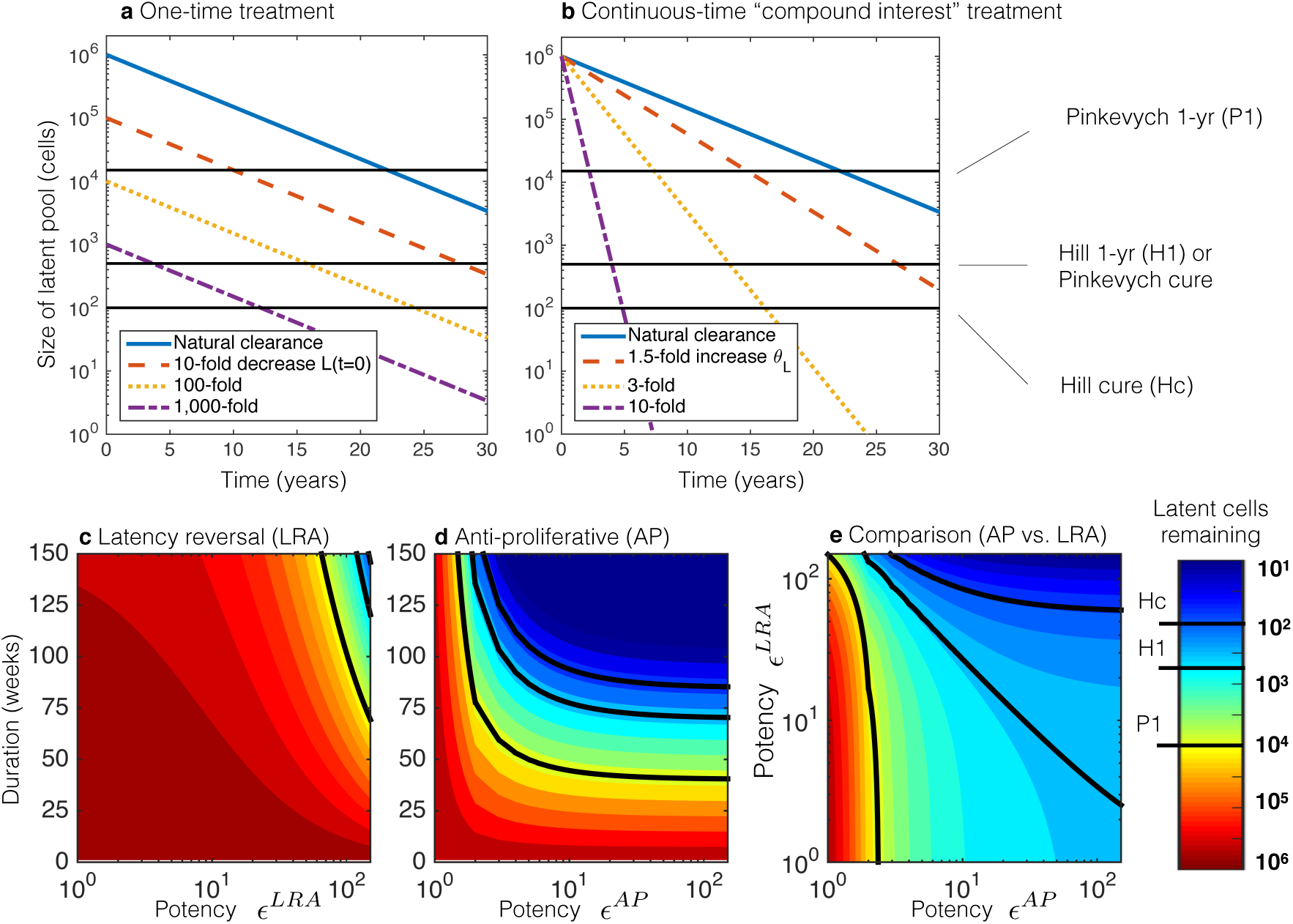
Simulated comparisons of latent reservoir eradication strategies on standard antiretroviral (ART) treatment. Treatment thresholds (discussed in Methods) are shown as solid black lines both in the plots and color bar, which is consistent between panels. **a**) One-time therapeutic reductions of the latent pool (*L*_0_). **b**) Continuous therapeutic increases in the clearance rate (*θ*_*L*_). Relatively small decreases in the clearance rate *θ*_*L*_ produce markedly faster times to cure than much larger decreases in the initial reservoir size. **c-e**) Latency reversal agent (LRA) and anti-proliferative (AP) therapies are given continuously for durations of weeks with potencies given in fold increase in activation rate (*ϵ*^*LRA*^) and fold decrease in proliferation rate (*ϵ*^*AP*^), respectively. The color bar is consistent between panels, and thresholds of cure are shown as solid black lines both on plots and on the color bar. **c**) Latency reversing agent therapy (LRA) administered alone requires years and potencies above 100 to achieve the cure thresholds. **d**) Anti-proliferative therapies (AP) administered alone lead to cure thresholds in 1-2 years provided potency is greater than 2-3. **e**) LRA and AP therapies are administered concurrently, and the reduction in the latent pool is measured at 70 weeks. Because the proliferation rate is naturally greater than the activation rate, increasing the AP potency has a much stronger effect than increasing the LRA potency.

Continuous-time interventions are more promising. Relatively small changes in *θ*_*L*_ in equation (4) lead to significant changes in the time to cure (Fig. 2b). On ART alone, estimated cure occurs at roughly 70 years [1]. However, just a 3-fold increase in clearance rate achieves Hill cure in fewer than 20 years. A 10-fold sustained increase requires only five years for Hill cure.

Further, when continuous-time therapies are given, outcomes improve more by extending duration than by equivalent increases in potency (Fig. 2c and 2d demonstrate this given the substantial asymmetry of the contours over their *y* = *x* axes). Analogous to the so-called “miracle of compound interest,” increasing the clearance rate for an extended duration produces profound latency reduction.

### Smaller reductions in proliferation rate achieve more rapid reservoir depletion than comparable relative increases in activation rate

Latency reversing therapy can be modeled with equation (3) if treatment is assumed to be a continuous-time multiplication of activation. Simulations at various potencies and therapy durations indicate both Hill and Pinkevych cure thresholds require more than a 100-fold multiplication of *ξ* sustained for two or three years, respectively (Fig. 2c).

The latent cell proliferation rate is considerably larger than the activation rate (*α*_*L*_ *» ξ*, Table 1). Thus, anti-proliferative therapies would clear the reservoir faster than equivalently potentlatency reversing strategies. When the reservoir of CD4^+^ T cells harboring replication-competent HIV is assumed to consist only of central memory cells (T_cm_), a 10-fold reduction in *α*_*cm*_ leads to Pinkevych 1-yr, Hill 1-yr, Pinkevych cure, and Hill cure in 0.8, 1.6, 1.6, and 1.8 years, respectively (Fig. 2d).

The improvement in cure time (when compared to an equivalent 10-fold increase in net reservoir clearance rate *θ*_*L*_) is possible because decreasing the proliferation rate means the net clearance rate approaches the latent cell death rate *δ*_*L*_. In fact, potency is relatively unimportant beyond reducing the proliferation rate by a factor of ten because the underlying death rate *δ*_*L*_ is the bound on clearance rate. The relative impact of anti-proliferative therapy is greater than that of latency reversing therapy when the two therapies are given concurrently for 70 weeks (Fig. 2e).

### Heterogeneity in reservoir cell types may necessitate prolonged anti-proliferative therapy

Recent studies indicate that the reservoir is heterogeneous, consisting of CD4^+^ central memory (T_cm_), naïve (T_n_), effector memory (T_em_), and stem cell-like memory (T_scm_) T cells. Further, reservoir cell composition differs dramatically among patients [12, 6, 23]. This heterogeneity suggests the potential for variable responses to anti-proliferative agents. Proliferation rates of T_cm_ (once per 66 days) exceed T_n_ (once every 500 days) but lag behind T_em_ proliferation rates (once every 21 days, Table 1). In our model T_scm_ are assumed to proliferate at the same frequency as T_n_ based on similar properties. We simulate possible reservoir profiles with different percentages of T_n_, T_cm_, and T_em_ in Fig. 3a-c. At least 7 years of treatment is needed for Pinkevych functional cure (Hill 1-yr) if slowly proliferating cells (T_n_ and/or T_scm_) comprise more than 20% of the reservoir. In contrast, an increased proportion of T_em_ has no clinically meaningful impact on time to cure. Slowly proliferating cells are predicted to comprise the entirety of the reservoir within two years of 10-fold anti-proliferative treatment regardless of initial percentage of T_n_ or T_scm_ (Fig. 3d-e).

**Figure 3:**
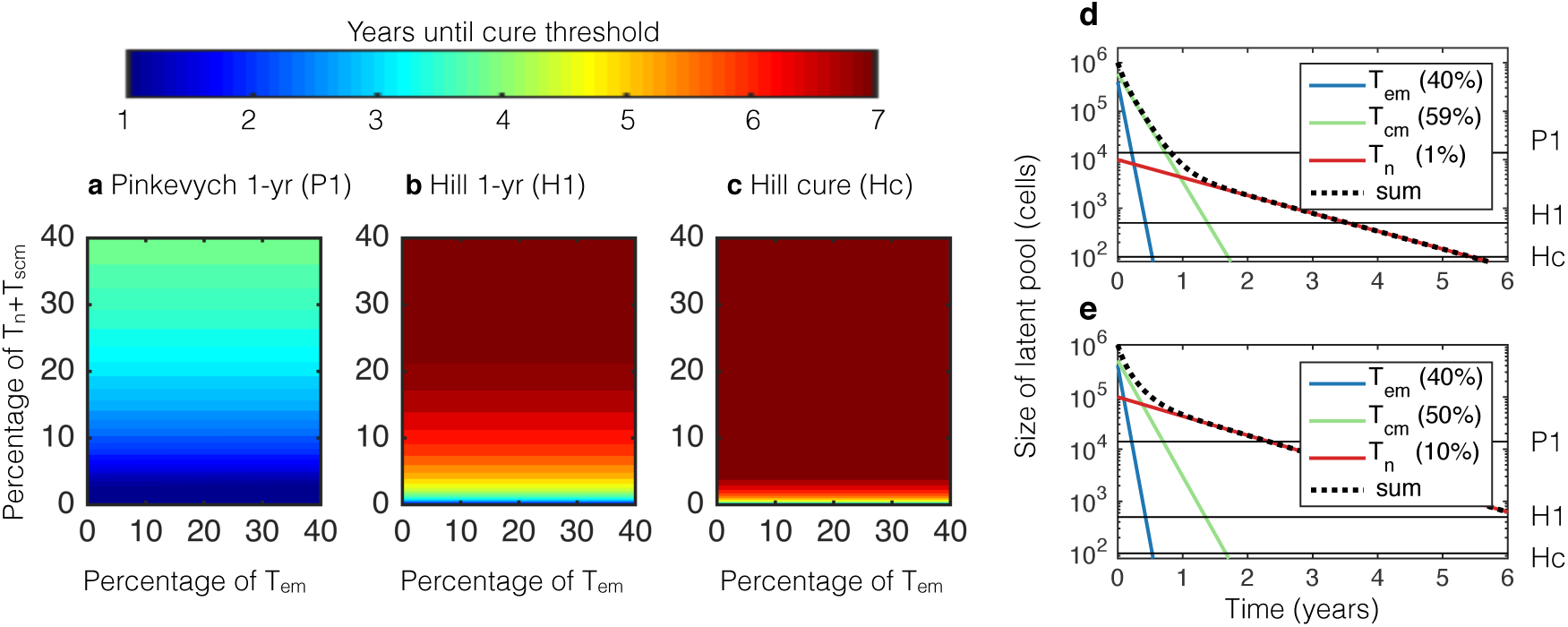
Simulated comparisons of anti-proliferative therapies on standard antiretroviral therapy (ART) assuming variable reservoir composition. Proliferation and death rates in Table 1. The potency of the therapy is *ϵ*^*AP*^ = 10 (*i.e.*, each cell type *i* has proliferation rate equal to *α*_*i*_/10 with *i* ∈ [em, cm, n]). Plausible initial compositions of the reservoir (*L*_*i*_(0)) are taken from experimental measurements [12, 6, 23]. It is assumed that the HIV activation rate *ξ* is equivalent across all reservoir subsets. **a-c**) Plots of times to therapeutic landmarks on long-term ART and anti-proliferative therapy with heterogeneous reservoir compositions consisting of effector memory (T_em_), central memory (T_cm_), and naïve plus stem cell-like memory (T_n_+T_scm_) CD4^+^ T cells. T_em_ and T_n_+T_scm_ percentages are shown with the remaining cells representing T_cm_. Times to one-year remission and functional cure are extremely sensitive to percentage of T_n_+T_scm_ but not percentage of T_em_. **d-e**) Continuous 10-fold therapeutic decreases in all proliferation rates (*α*_*i*_) result in Hill 1-yr in **d**) 3.5 years assuming T_n_+T_scm_=1% and **e**) 6 years assuming T_n_+T_scm_=10%. The reservoir is predicted to become T_n_+T_scm_ dominant within 2 years under both assumptions, providing an indicator to gauge the success of anti-proliferative therapy in potential experiments.

The uncertainty in the reservoir composition tempers the results in Fig. 2. On the other hand, our model assumes that the HIV activation rate *ξ* is equivalent across all CD4^+^ T cell reservoir subsets. It is biologically plausible, though unproven, that latent cell proliferation and activation are linked processes and that therefore HIV rarely or never activates from resting T_n_ or T_scm_. Under this assumption, functional cure might occur once T_em_ and T_cm_ reservoirs have been reduced to the Hill cure level, *i.e.* approximately 1.5 years in Fig. 3d and 3e.

### Initial reservoir size, anti-proliferative potency, and reservoir cell subtypes pre-dict time to cure

Using literature-derived ranges for the parameters of interest, we completed a global sensitivity analysis to examine which factors might impact time to cure in a heterogeneous patient pool developed by Latin Hypercube sampling of a broad parameter space [24] (Fig. 4). We correlate variables with time to cure on ART/anti-proliferative combination therapy. Varying the probability of latency given infection (*τ*) does not change time to cure. Similarly, varying the basic reproductive number on ART (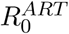), a measure of ART efficacy, defined as the number of new infected cells generated by one infected cell during ART, does not change time to cure. On the other hand, as the pre-treatment size of the latent pool *L*_0_ increases, the necessary time to cure also increases. Increasing anti-proliferative therapy potency *ϵ*^*AP*^ decreases cure time. Increasing percentages of naïve T cells *L*_*n*_(0)*/L*_0_ in the latent reservoir delay the time to cure while a faster latent decay rate *θ*_*L*_ hastens cure. Finally, we simulated the possibility of a diminishing impact of anti-proliferative therapy over time in Fig. 5. The simulation shows that when potency decreases by less than 5% per month, cure thresholds are still achieved within 10 years of ART and anti-proliferative treatment. The fastest waning of potency (20% per month) results in return to the natural clearance rate within the first 2 years of therapy prompting longer times to cure.

**Figure 5:**
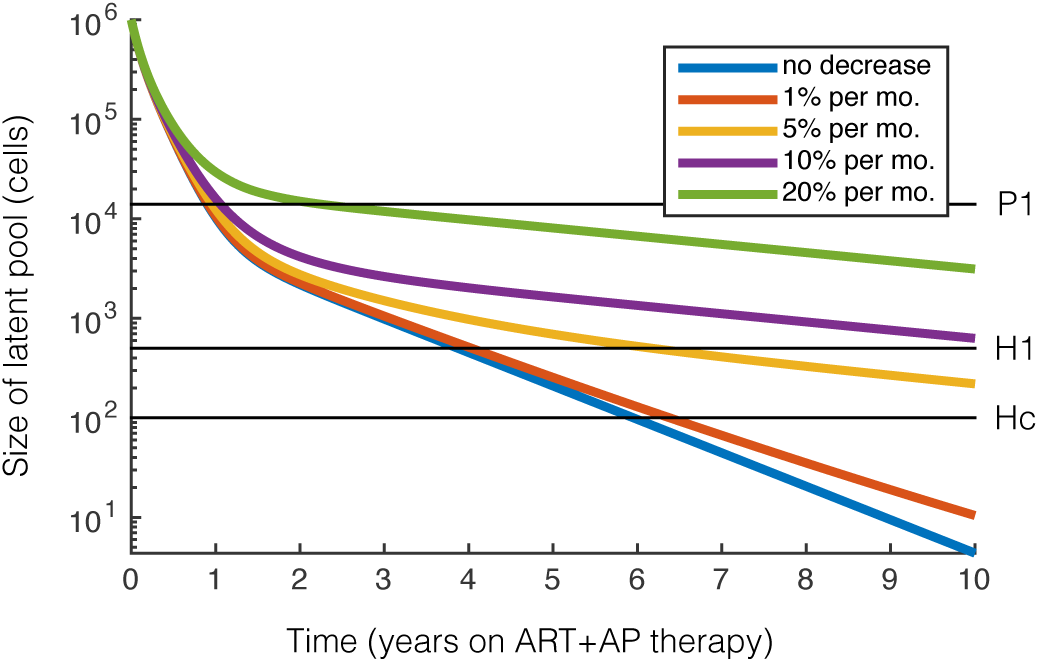
Waning anti-proliferative potency over-time modulates cure. Latent reservoir dynamics on combined ART and anti-proliferative therapy simulated for waning potency of anti-proliferative therapy over time. The latent reservoir size is shown with horizontal black lines corresponding to the cure threshholds used throughout the paper. Cure thresholds are achieved within 10 years if potency decreases by less than 5% per month considering 1% naïve T cells (*L*_*n*_(0)*/L*_0_ = 0.01) and initial anti-proliferative potency *ϵ*^*AP*^ = 5.

**Figure 4:**
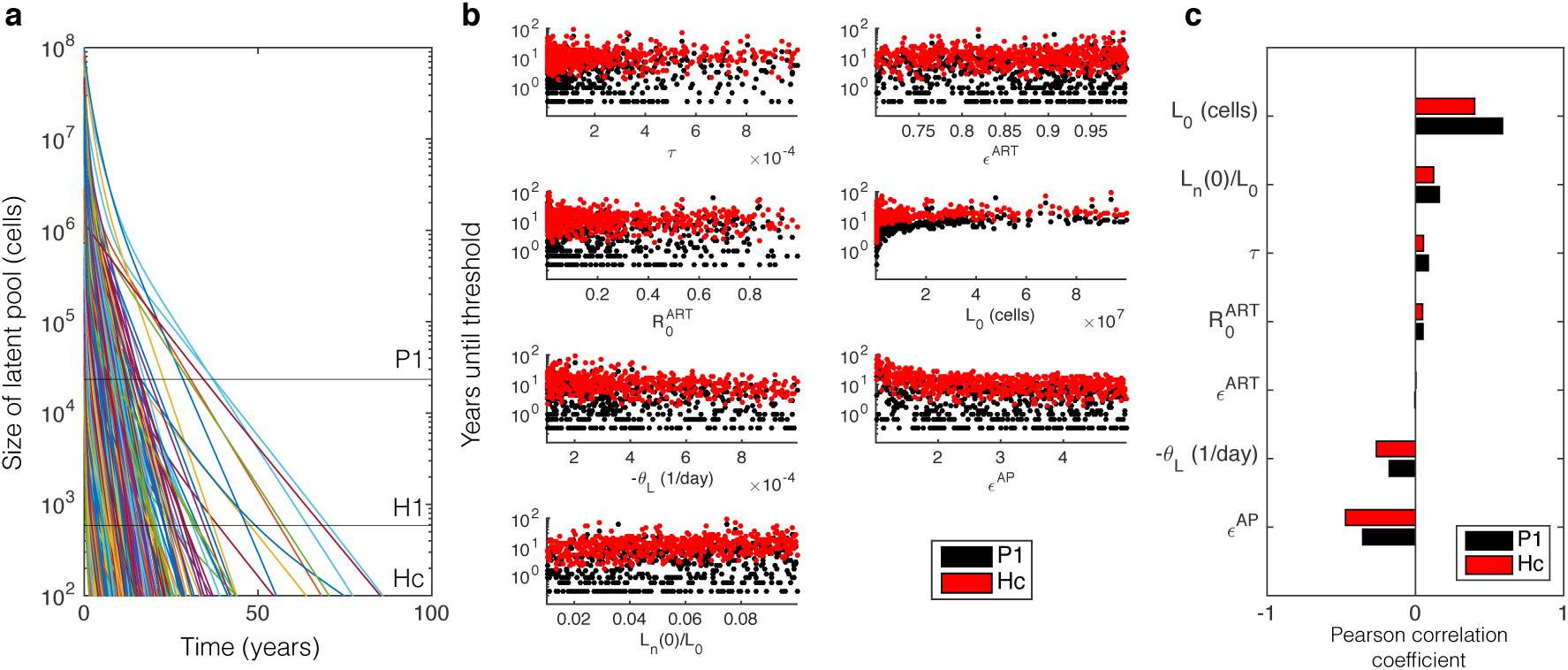
Global sensitivity analysis. We use the ranges of parameters from Supplementary Table S4 online. **a**) 1,000 simulations drawn from Latin Hypercube sample parameter sets where *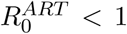* are shown to demonstrate the variability of latent pool dynamics with respect to all combinations of parameter ranges. **b**) The time until each cure threshold, Pinkevych 1-yr (P1) and Hill cure (Hc), are calculated as the time when the latent reservoir contains fewer than 20,000 and 200 cells respectively. In some cases cures are achieved within months. In others, cure requires many years. **c**) Pearson correlation coefficients indicate the correlations between each variable and time to cure. *L*_0_ is the initial number of latent cells. *L*_*n*_(0)*/L*_0_ is the initial fraction of naïve cells 0 in the latent pool. *τ* is the probability of latency given infection. *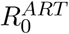* is the basic reproductive number on ART. *ϵ* ^*ART*^ is the percent decrease in viral infectivity in the presence of ART. *θ*_*L*_ is the decay rate of latent cells. *ϵ*^*AP*^ is the fold reduction in proliferation rate.

### Model output is congruent with available clinical data

Chapuis *et al.* treated eight ART-suppressed, HIV-infected patients with 24 weeks of mycophenolate mofetil, a licensed anti-proliferative agent. As a marker of anti-proliferative effect, the percentages of Ki67^+^CD4^+^ T cells were measured before and after MMF treatment (2 *×* 500 mg daily) and were found to have decreased on average 2.2-fold. Incorporating that reduction in latent cell proliferation rate *ϵ*^*AP*^ = 2.2 over 24 weeks of treatment, we estimate a 10- to 40-fold reduction in the latent reservoir (see Fig. 2d). Chapuis *et al.* found a 10- to 100-fold reduction in infectious units per million (IUPM) by quantitative viral outgrowth assay in five patients, comparable to our estimate [25]. These reductions far exceed natural reservoir clearance rates and are consistent with a therapeutic effect [26].

García *et al.* assessed the effect of MMF (2 *×* 250 mg daily) on HIV in the context of ART treatment interruption [27]. Seventeen HIV-infected patients received ART for a year and then were randomized into a control group that remained on ART only and an experimental group that also received MMF for 17 weeks. ART was interrupted in both groups and viral rebound assessed. MMF inhibited CD4^+^ T cell proliferation (as measured by an *in vitro* assay) in six of nine MMF recipients (responders). The time to rebound was 1-4 weeks in the control group and 6-12 weeks in the MMF-responder group. Using results from Pinkevych *et al.*, a median time to rebound of seven weeks (see Fig. 5b of Ref. [22]) corresponds to a 7-fold decrease in the latent reservoir. Using results from Hill *et al.*, the same median time to detection of seven weeks (see Fig. 4 of Ref. [21]) corresponds to a 50-fold reduction in the latent reservoir. These calculations are congruent with our model’s estimate that 17 weeks of MMF treatment at potency *ϵ*^*AP*^ = 2.2 leads to a 10-fold reduction in the reservoir.

**Table 4:** Viral burst and clearance rates. Note: [56] used the data from [60] but used the geometric mean rather than the arithmetic mean, *i.e.* 18.8 rather than 23.

| Parameter | Perelson [45] | Huang [58] | Luo [56] | Units |
| --- | --- | --- | --- | --- |
| $\pi$ | 1200 | 976 | $5.9 \times 10^3$ | copies/cell-day |
| $\gamma$ | 2.4 | 3.06 | 18.8 | per day |
| $\pi/\gamma$ | 500 | 319 | 314 | copies/cell |

### MMF decreases proliferation in CEM cells, CD4^+^ T cells from HIV positive and negative donors, and all CD4^+^ T cell subsets

To explain the heterogeneous impact of MMF treatment (three of six in Chapuis *et al.* did not demonstrate a meaningful reservoir clearance; three of nine patients in García *et al.* had a weak anti-proliferative response to MMF and no delay in HIV rebound upon ART cessation), we conducted an *in vitro* study of MMF pharmacodynamics. We titrated the capacity of mycophenolic acid to inhibit spontaneous proliferation of cells from a human T lymphoblastoid cell line (CEM cells)[28] and identified a steep Hill slope of -3.7 (Fig. 6a). A Hill slope with absolute value greater than one indicates cooperative binding at the site of drug action and implies a sharp transition from negligible to complete therapeutic effect at a specific drug concentration. These results explain how patients with inadequate MMF dosage could have a limited anti-proliferative effect.

**Figure 6:**
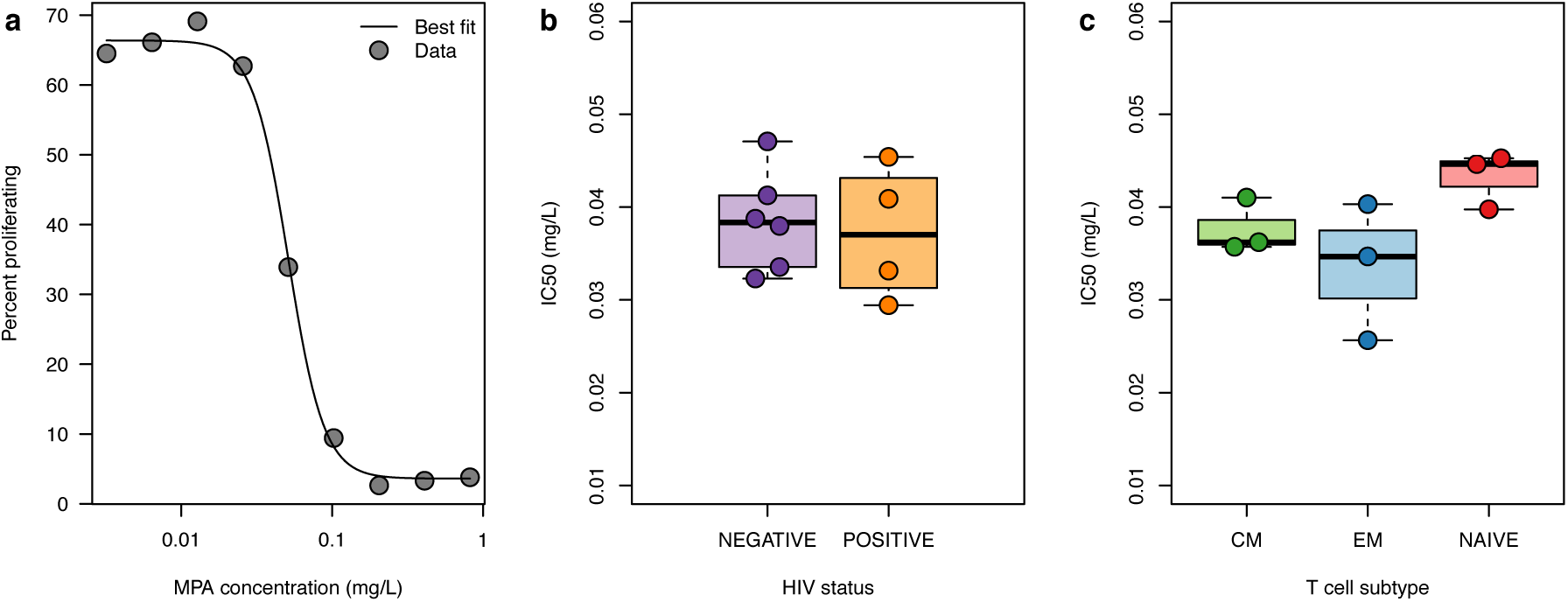
MMF pharmacodynamics. Pure mycophenolic acid (MPA) was added to CEM cells at varying concentrations and proliferation of CEM cells was measured to determine a dose-response curve and Hill slope. CD4^+^ T cells from stored peripheral blood mononuclear cell samples from 10 participants (4 HIV-infected, 6 HIV-uninfected) were stimulated to proliferate. CD4^+^ T cells from 3 HIV-negative subjects were sorted into effector memory (EM), central memory (CM), and naïve subsets. Pure MPA was added to these cells at varying concentrations in order to determine IC50s for MPA. **a**) Dose-response curve with percentages of CEM cells proliferating at varying doses of MPA. The Hill slope is -3.7. **b**) 4 samples from HIV-positive participants and 6 samples from HIV-negative participants had similar IC50s. **c**) IC50s were similar among CD4^+^ effector memory (EM), central memory (CM), and naïve T cell subsets.

**Table 3:** Infectivity.

| Parameter | Perelson [45] | Huang [58] | Luo [56] | Units |
| --- | --- | --- | --- | --- |
| $\beta$ | $2.4 \times 10^{-5}$ | $1.7 \times 10^{-5}$ | $3.9 \times 10^{-3}$ | $\mu\text{L per day}$ |

We tested the capacity of mycophenolic acid (MPA), the active metabolite of MMF, to inhibit CD4^+^ T cell proliferation in CD4^+^ T cells from four HIV-positive and six HIV-negative participants and found similar IC50s (Fig. 6b). Further, CD4^+^ T cells from three HIV-negative participants were sorted into central-memory, effector-memory, and naïve subsets. Similar proliferation inhibition was observed in all three cell subsets (Fig. 6c). These results suggest a potential for MMF to deplete the HIV reservoir.

## Discussion

We developed a mathematical model of HIV dynamics to study various cure strategies [21, 29]. We demonstrate that minor reductions in CD4^+^ T cell proliferation rates would exhibit powerful reductions in the latent reservoir when therapy duration is extended over time. We call this proposed strategy “compound interest cure” due to the correspondence with financial modeling.

Our results are relevant because the HIV cure strategy most rigorously being tested in humans— latency reversal therapy (“shock-and-kill”)—may not capitalize on the advantages of a compound interest approach. Promising latency reversing agents are typically dosed over short time-frames due to concerns about toxicity. T cell activation does not always lead to induction of HIV replication providing another potential limitation of latency reversing therapy [30]. Furthermore, even if these agents exert a large relative impact on the activation rate of memory CD4^+^ T cells, we predict the reduction in the reservoir may be insignificant given that the natural activation rate is orders of magnitude lower than proliferation and death rates. Latency reversal agents are the being considered in conjunction with other interventions such as engineered antibodies and/or T cells. These combined approaches carry additional unknown toxicities and rely on the effectiveness of latency reversal agents. Most challenging of all, these experimental therapies could be prohibitively expensive to implement globally.

The theoretical potential of the anti-proliferative approach is worthy of a clinical trial given the existence of licensed medications that limit T cell proliferation, including MMF. In line with our prediction that duration is more important than potency, these drugs are dosed over months to years for rheumatologic diseases and preventing rejection after solid organ transplant. The most frequent side effects reported are gastrointestinal symptoms and increased risk of infection though the latter risk is obscured by concurrent use of high-dose glucocorticoids [31]. MMF has been given to several hundred HIV-infected patients suppressed on ART[32, 33, 34, 35, 36, 37, 38, 39, 27, 25, 40] (reviewed in **Supplementary information online**). In this population, neither opportunistic infections nor adverse events were increased, and CD4^+^ T cell counts did not decrease significantly during therapy. We hypothesize that whereas MMF decreases proliferation of existing CD4^+^ T cells, it does not suppress thymic replenishment of these cells. Finally, MMF did not counteract the effects of ART [27, 25], and we do not expect viral drug resistance or ongoing viral evolution to occur on anti-proliferative therapy. Despite these reassuring findings, future studies of HIV-infected patients on anti-proliferative agents will require extremely close monitoring for drug toxicity and immunosuppression. In addition, mycophenolic acid has a large Hill coefficient, suggesting a narrow therapeutic range. We suspect that the participants who did not respond to MMF in the clinical studies described above [25, 27] required higher drug concentrations.

Our model suggests that slowly proliferating cells in the reservoir could present a barrier to rapid eradication of latently HIV-infected cells. Therefore, anti-proliferative strategies may face a challenge akin to the cancer stem cell paradox, whereby only the rapidly proliferating tumor cells are quickly expunged with chemotherapy. For example, tyrosine kinase inhibitors suppress proliferation of cancer cells in chronic myelogenous leukemia (CML). While many patients achieve “undetectable minimal residual disease,” some patients relapse to pre-therapy levels of disease following therapy cessation—perhaps due to slowly proliferating residual cancer stem cells [41]. Additional limitations could include insufficient anti-proliferative drug delivery to anatomic sanctuaries, certain cellular subsets that are unaffected by treatment, and cytokine-driven feedback mechanisms that compensate for decreased proliferation by increasing memory CD4^+^ T cell lifespan. These challenges might be countered by combining anti-proliferative agents with other cure therapies. Avoidance of nucleoside and nucleotide reverse transcriptase inhibitors, which may enhance T cell proliferation, could provide an important adjunctive benefit [42, 43].

The anti-proliferative approach is attractive because it is readily testable without the considerable research and development expenditures required for other HIV cure strategies. Anti-proliferative approaches require minimal potency relative to latency reversing agents, and T cell anti-proliferative medications are well studied mainstays of organ rejection prevention. Therefore, we propose trials with anti-proliferative agents as an important next step in the HIV cure agenda.

## Methods

### Latent reservoir dynamic model

We based our model (schematic in Fig. 1) on previous HIV dynamics models [29, 44]. We follow the concentrations [cells/*μ*L] of susceptible CD4^+^ T cells *S*, latently infected cells *L*, actively infected cells *A*, and plasma viral load *V* [copies/*μ*L] over time. The system of ordinary differential equations (using the over-dot to denote derivative in time)

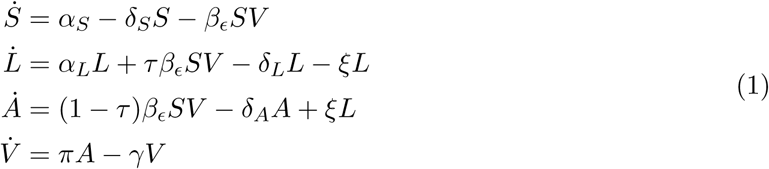

tracks these state variables. We define *α*_*S*_ [cells/*μ*L-day] as the constant growth rate of susceptible cells, *δ*_*S*_ [1/day] as the death rate of susceptible cells, and *β*_*ϵ*_ = (1 *-ϵ*)*β* [*μ*L/virus-day] as the therapy-dependent infectivity. We define *ϵ* [unitless] as the ART efficacy, ranging from 0 (meaning no therapy) to 1 (meaning perfect therapy). *α*_*L*_ and *δ*_*L*_ [1/day] are the proliferation and death rates of latent cells, respectively. The death rate of actively infected cells is δ_*A*_, and the proliferation rate of activated cells *α*_*A*_ *≈* 0 is likely negligible [45]. *τ* [unitless] is the probability of latency given infection, and *ξ* [1/day] is the rate of transition from latent to actively infected cells. The viral production rate is *π* [virions/cell-day], which describes the aggregate rate of constant viral leakage and burst upon cell death. *γ* [1/day] is the HIV clearance rate. Parameter values are given in Table 1.

Additional calculations including derivations of equilibrium solutions and stability analysis as well as further discussion of model parameter derivations are presented in the **Supplementary information online**.

### The compound interest formula

In the **Supplementary information online**, we determine the critical drug efficacy *ϵ*_*c*_, the value of *ϵ* above which viral load quickly decays. Moreover, when *ϵ > ϵ* _*c*_, we can consider the latent cell equation in isolation:

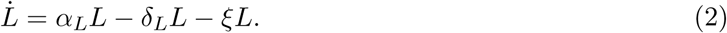

Defining the initial number of latent cells as *L*_0_ gives

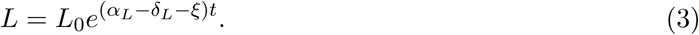

Equation (3) implies that the clearance rate of latently infected cells is a function of their proliferation, death, and activation rates. Defining the total clearance rate *θ*_*L*_ = *α*_*L*_ - *δ*_*L*_ - *ξ*, we see a mathematical correspondence to the principle of continuous compound interest with *L*_0_ as the principal investment and *θ*_*L*_ as the interest rate:

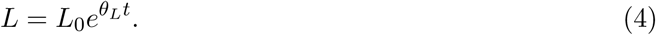

Experimental measurements indicate an average latent cell half-life of 44 months (*θ*_*L*_ = −5.2 *×* 10^-4^ per day) [1, 26] and an average latent reservoir size *L*_0_ of one-million cells[1]. Note that when *θ*_*L*_ *<* 0, the latent reservoir is cleared exponentially. Alternatively, if *α*_*L*_ exceeds the sum of *ξ* and *δ*_*L*_, *L* grows indefinitely.

### Composition of the latent reservoir: modeling T cell subsets

We include heterogeneity in T cell phenotype into the model by splitting the differential equation for the latent cells into three differential equations, one for each subtype *L*_*i*_ with *i* ∈ [*cm, em, n*]. We ignore transitions between phenotype because the composition of the reservoir is reasonably stable over time [12]. Our extended model is the system

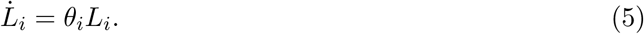

The total number of latent cells is the sum of the subset populations, *L* = ∑_*i*_ *L*_*i*_, and solution is

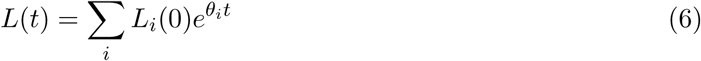

where *θ*_*i*_ = *α*_*i*_ - *δ*_*i*_ - *ξ*, and *L*_*i*_(0) are the initial numbers of each subtype.

Simulations assume the same net clearance rate and activation rates among subsets, but different proliferation rates *α*_*i*_ and different calculated death rates *δ*_*i*_ = *α*_*i*_ - *θ*_*L*_ - *ξ*. The initial conditions for each subtype *L*_*i*_(0) are inclusive of several varying measurements in the literature [12, 6, 23]. We consider T_tm_ to have the same proliferation rates as T_cm_. Similarly, we characterize stem-cell-like memory CD4^+^ T cells (T_scm_) as T_n_ given their slow turnover rate. Of note, these are conservative estimates that would not favor anti-proliferative therapy. In Fig. 5, we allow the anti-proliferative potency to decrease over time by assuming *α*_*i*_(*t*) = *α*_*i*_[1 + (*ϵ*^*AP*^ - 1) exp(-*φt*)] for each T cell subset in Eq. 5. Here *φ* is the waning potency rate that ranges from 0–20% per month. We assume the initial potency is a 5-fold decrease *ϵ*^*AP*^ = 5, and we use a 1 million cell reservoir having 1% naïve T cells. We solve the equation for each subset numerically with ode23s in Matlab, summing the subset dynamics after solving.

### Reservoir reduction targets for cure strategies

We use experimentally derived thresholds to compare potential cure therapies in the framework of our model. Hill *et al.* employed a stochastic model to estimate that a 2,000-fold reduction in the latent pool would result in HIV suppression off ART for a median of one year. After a 10,000-fold reduction in latent cells, 50% of patients would remain functionally cured (ART-free remission for at least 30 years) [21]. Pinkevych *et al.* inferred from analytic treatment interruption data that decreasing the latent reservoir 50-70-fold would lead to HIV remission in 50% of patients for one year [22]. Using the Pinkevych *et al.* results, we extrapolate a functional cure threshold as a 2,500-fold decrease in the reservoir size (**Supplementary information online**). Given ongoing debate in the field, we consider all four thresholds—henceforth referred to as Hill 1-yr, Hill cure, Pinkevych 1-yr, and Pinkevych cure.

### Sensitivity analysis

To examine the full range of possible outcomes we completed a global sensitivity analysis of the model in which all variables were simultaneously varied in the ranges of Table S5 by logarithmically covering Latin Hypercube sampling [24]. The simulations were carried out in Matlab using lhsdesign and ode23s. We correlated each parameter of interest with the time to reach the Hill and Pinkevych cure thresholds. Calling the time-to-cure *T*, correlations were calculated with the Pearson correlation coefficient: the covariance of *T* with each parameter of interest *p* normalized by both the standard deviation of *T* and that of *p*, that is *ρ* = cov(*T, p*)*/σ*_*T*_ *σ*_*p*_. 1,000 simulations were carried out, keeping only the parameter combinations leading to reservoir decay, i.e. those satisfying 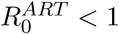.

### Mycophenolic acid anti-proliferation assay methods

Blood samples for the MPA *in vitro* studies were obtained from ART-treated, HIV-infected and healthy, HIV-negative men at the HIV Vaccine Trials Unit Clinic in Seattle, Washington. All procedures were approved by the Institutional Review Boards of the University of Washington and the Fred Hutchinson Cancer Research Center (IRB 1830 and 5567) and were performed in accordance with institutional guidelines and regulations. Written informed consent was obtained from each donor.

Cells were labeled using the CellTrace Violet Cell Proliferation Kit (Invitrogen) by incubation in 40 *μ*M CellTrace Violet in Roswell Park Memorial Institute (RPMI) cell culture media with penicillin/streptomycin and L-glutamine (Gibco) plus 10% fetal bovine serum (Gemini Bio-Products)(R-10 media) for five minutes at room temperature [46] followed by washing twice with R-10. Peripheral blood mononuclear cells were stimulated with 1 *μ*g/mL staphylococcal enterotoxin B (SEB; Sigma-Aldrich) and 10 IU/mL IL-2 (Peprotech). Sorted CD4+ T cell subsets (naïve, effector memory, and central memory) were stimulated with Dynabeads Human T-Activator CD3/CD28 beads (Gibco) at a bead to cell ratio of 1:1 with 10 IU/mL IL-2. CEM cells were not stimulated, as they proliferate continuously. Pure mycophenolic acid (Sigma-Aldrich), the active metabolite of MMF, was added at concentrations ranging from 0.01 to 2.56 *μ*M. Cells were cultured in R-10 for 72 h.

After the culture period, cells were washed and stained with Fixable Live/Dead Yellow (Invitrogen), followed by CD45RA FITC, CD4 PE-Cy5, CCR7 BV785 (all BD), and CD3 ECD (Beckman Coulter) at the minimum saturating doses. Cells were then fixed with 1% paraformaldehyde and acquired on a five-laser BD LSRII flow cytometer (355, 405, 488, 535, and 633 nm). Live, single CD4+ T cells were gated into “proliferated” or “not proliferated” on the basis of CellTrace Violet fluorescence.

The IC50s and Hill slope were calculated using the drc package in R (**Supplementary information online**)[47, 48].

### Appendix I. Additional modeling methods

#### Equilibrium solutions leading to decoupled equations for latently infected cells

Our model for HIV including latently infected cells and antiretroviral therapy (ART) is described completely in the main body with all rates specified in Table 1. Here we present the ordinary differential equations (ODEs) without further clarification:

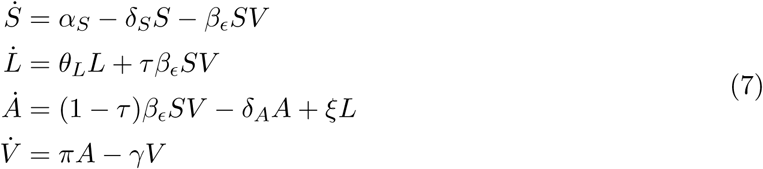

Equilibrium solutions (denoted by the asterisk) can be calculated by setting 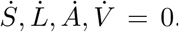. The system has two equilibrium solutions:

1. “Viral-free equilibrium”, the values at which the system rests prior to infection

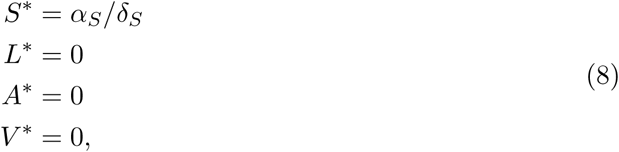
2. “Setpoint equilibrium”, the values at which the system has durable viral infection: commonly referred to as “viral setpoint” in clinical HIV care.

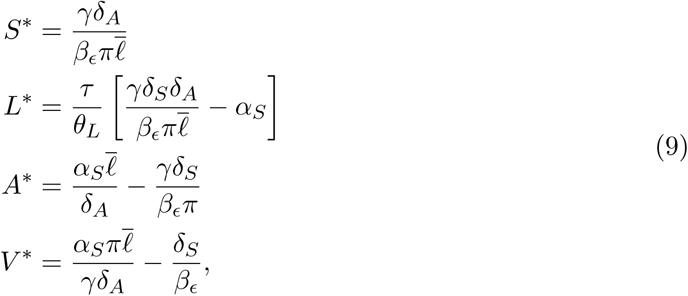

where 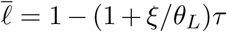. We use the 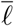 notation to simplify the appearance of the equations and choose the lower-case 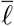, as this factor encapsulates all the latent dynamics. For *τ «* 1, we have *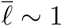*, and *τ* is far less than one in the literature (see Table 5).

**Table 5:** Fraction of infected cells that enter a latent state.

| Parameter | Callaway [15] | Jones [64] | Conway [29] | Units |
| --- | --- | --- | --- | --- |
| $\tau$ | $10^{-6}$ | $10^{-3}$ | $10^{-4}$ | unitless |

#### Stability analysis of equilibrium solutions: Calculating the basic reproductive number confirms that decoupling latent cell equations is valid when ART efficacy is above the critical value

By linearizing our system of equations, we address the local stability of the equilibrium points. We begin by defining our state variables as a vector **x** = [*S, L, A, V*]^*T*^ such that we can express the system of ODEs as **F**(**x**) = *∂*_*t*_**x**. Then, we Taylor expand this function around the equilibrium point **x**\* where **F**(**x**\*) = 0:

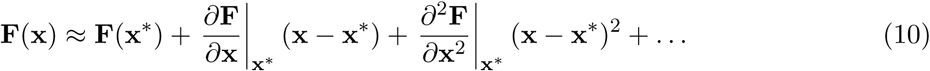

By construction the first term in the expansion is zero. Calling a small Δ**x** *«* 1 = **x** - **x**\*, we can neglect terms of *𝒪*(Δ**x**^2^) and higher. Using **x** = **x**\* + ∆**x** we can rewrite

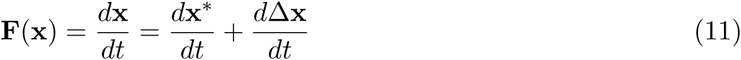

and because **x**\* is not a function of time we are left with the linear equation we desire:

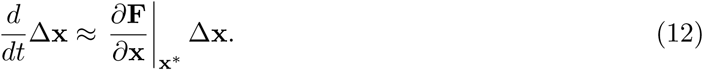

The matrix **J** = *∂***_x_F** is referred to as the Jacobian matrix for our model, and is in complete form:

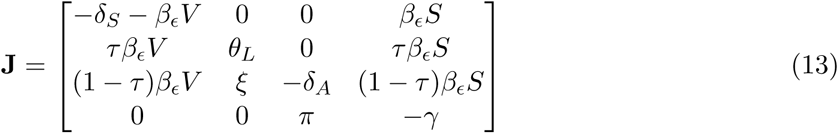

We can evaluate the Jacobian at both equilibria derived above (Eq. 8 and Eq. 9). The eigenvalues of the Jacobian then govern how perturbations near equilibrium behave. If all eigenvalues *λ*_*j*_ of the Jacobian have negative real components Re(*λ*_*j*_) *<* 0, perturbations decay back to equilibrium and the equilibrium is deemed stable [49]. Before infection, we input the equilibrium solutions (Eq. 8) and the Jacobian of the viral free equilibrium (subscript vfe) is

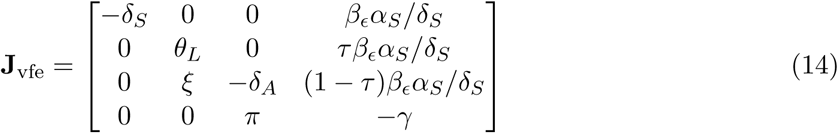

We can calculate the eigenvalues of this rather sparse matrix. Taking the determinant of 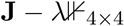 and setting this equal to zero we find

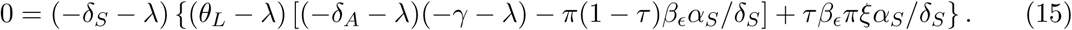

We immediately identify the eigenvalue *λ*_1_ *≈ -δ*_*S*_. Then, noting that the trailing term *τβ* _*ϵ*_ *πξα*_*S*_/*δ*_*S*_ is many orders of magnitude smaller than the other quantities (see parameter values in Table 1 of main body), we drop this term and identify a second eigenvalue *λ*_2_ *≈ θ*_*L*_. We remind the reader that *θ*_*L*_ = *α*_*L*_ - *δ*_*L*_ - *ξ* is a negative number and defined as the net clearance rate of the latent reservoir per Ref. [1]. Thus, we have identified two negative eigenvalues, meaning that perturbations to the viral free equilibrium return to equilibrium along those respective eigendirections. Solving for the remaining eigenvalues using the expression within the square brackets leads to

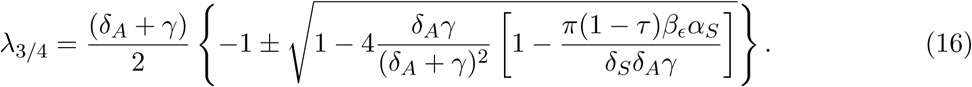

We define the quantity

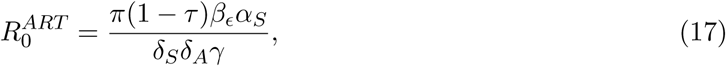

calling it the basic reproductive number that depends on the ART drug efficacy *ϵ*. By following through calculations (noting bounds on the factor 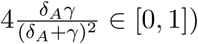) it can be shown that if 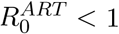 both *λ*_*3/4*_ *<* 0 such that the viral free equilibrium is stable. Letting *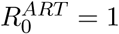* makes *λ*_3_ = 0 and *λ*_4_ *<* 0, while *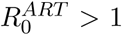* results in *λ*_3_ *≥* 0 while *λ*_4_ *<* 0. The basic reproductive number controls the stability of the viral free equilibrium, only when *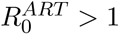* will an infection take off. Note that the same result can be verified using the next-generation matrix method [50]

To calculate the “critical drug efficacy” *ϵ*_*c*_, we solve for the drug efficacy that makes *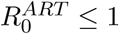*. This yields

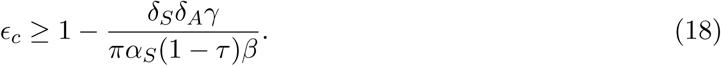

or using the typical definition of the basic HIV dynamics model 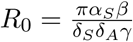, we see that

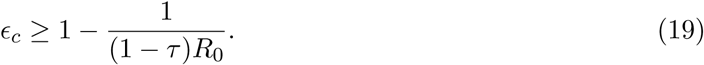

For our parameter values, *R*_0_ *∼* 8 as expected for HIV, guaranteeing instability of the viral free equilibrium for any introduction of virus.

Observations from patient data show that primary infections in humans lead to a durable viral setpoint. The Jacobian of our model evaluated at the endemic equilibrium can be written by inserting the equations Eq. 8 into Eq. 13. By calculating the eigenvalues of this Jacobian, we assess the stability of the setpoint equilibrium. Numerical computation of the eigenvalues of this matrix (using `eig()` in Matlab) for varying values of the drug efficacy are presented in Fig. 7. Here we find that above the critical drug efficacy *ϵ*_*c*_ (for our parameters *ϵ*_*c*_ *∼* 85%), the real part of the third eigenvalue becomes positive. Therefore, while the setpoint equilibrium is stable for low drug levels, *ϵ < ϵ*_*c*_, above the critical efficacy the setpoint is no longer stable. At precisely this drug level the viral free equilibrium becomes stable again, which can be calculated numerically, or seen from the analytical derivation above.

**Figure 7:**
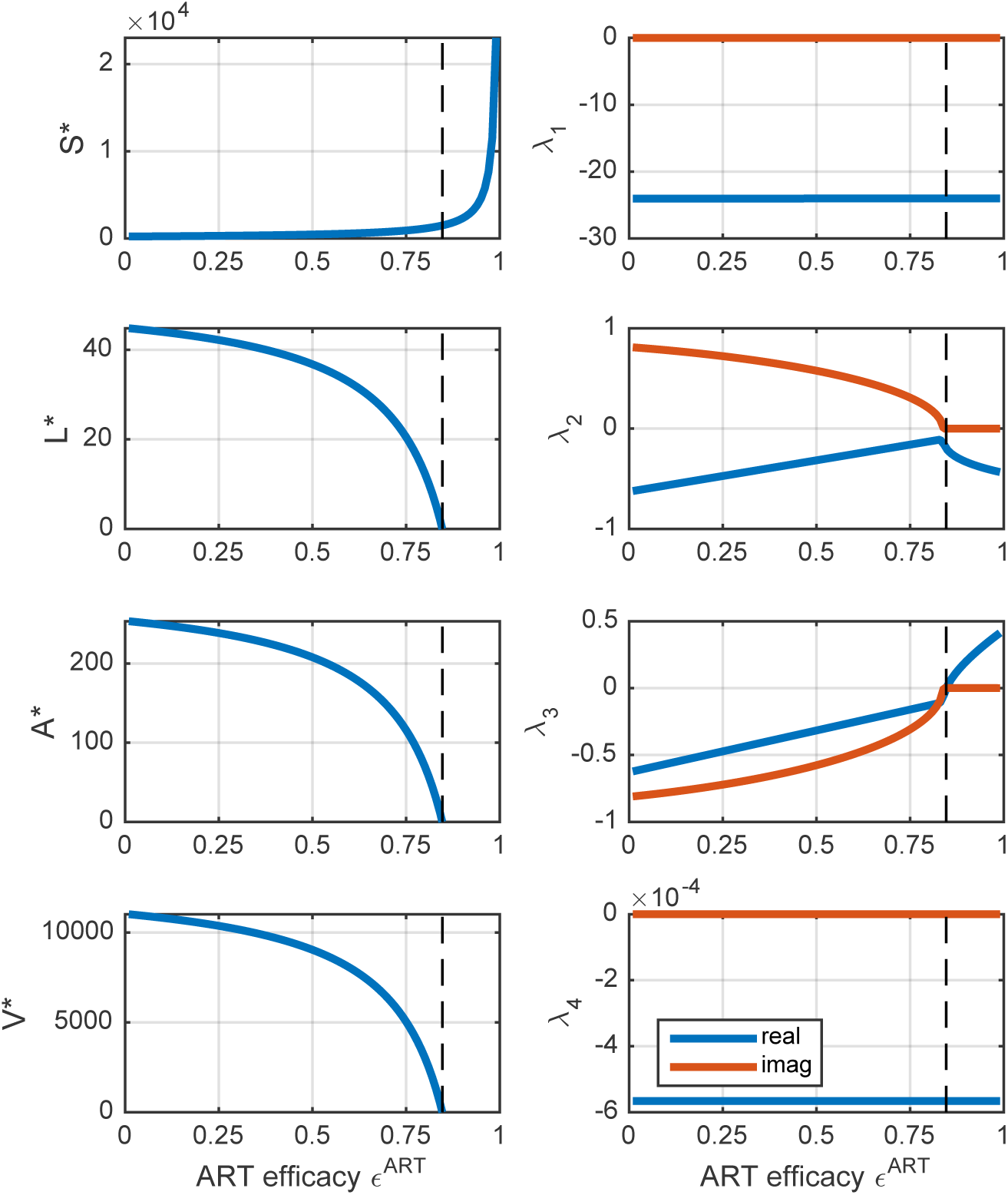
Setpoint stability depends on the efficacy of antiretroviral therapy (ART). Plots of the equilibrium concentrations and the eigenvalues of **J** (Eq. 13) evaluated at those viral setpoint equilibrium (Eq. 9) values at varying ART efficacy *ϵ*^*ART*^. The critical therapy thresholds are illustrated by the vertical dashed line (with our parameters and Eq. 18 we can calculate *ϵ*_*c*_ *∼* 85%). At this value the real part of the third eigenvalue becomes positive making the viral equilibrium unstable.

Furthermore, calculating the eigenvalues of the viral free equilibrium shows that above the critical efficacy, the viral free equilibrium becomes stable (all eigenvalues having negative real parts). For our parameter range, the eigenvalues (-0.2, -0.0006, -0.3, -24) approximate the death or clearance rates of susceptible cells, latent cells, active cells, and virus respectively. This result agrees with our analytical approximation of the first 2 eigenvalues above. Only the eigenvalue related to the active cell rate is noticeably increased relative to its natural rate, presumably due to coupling between two eigendirections. Most importantly, the disparate timescales allow adiabatic decoupling of the processes [51]. That is, because the solution for the return to equilibrium corresponds to the solution of

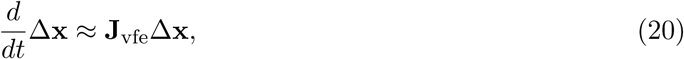

or using our eigenvalue decomposition, the complete solution to the dynamics of a perturbation from the viral free equilibrium can be expressed as

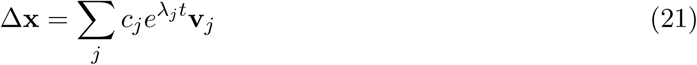

where **v**_*j*_ are the 4 eigendirections, not explicitly written here. The timescale for decay of each eigendirection is proportional to 1/*λ*_*j*_. For example, the fastest eigendirection *λ*_4_ becomes negligible in the timescale of a day, corresponding to viral clearance. Two of the other eigenvalues have timescales of around 1 week, corresponding to clearance of active cells. However, the time required to completely return to viral free equilibrium (*t*\*) depends on the slowest rate or the smallest magnitude eigenvalue as *t*\* *≈* 1/ min *|λ*_*j*_*|* [49]. In our case, the smallest eigenvalue is approximately *θ*_*L*_ so that complete return to viral free equilibrium ultimately depends only on the latent cell clearance rate. This justifies our approximation that virus is negligible soon after ART initiation and that the model for long term ART can be simplified to which has the solution

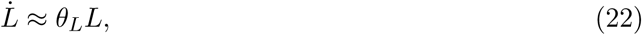

which has the solution

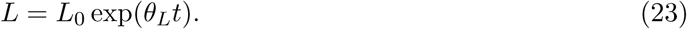

In summary, our model system rests at the viral free equilibrium prior to infection. Any virus results in HIV infection which progresses and settles at a stable viral setpoint equilibrium. However, ART disrupts this equilibrium, making it unstable and driving the system to return to viral free equilibrium. This movement toward viral-free equilibrium is slow and is limited by the rate of latent cell decay. But, this limiting decay is much slower than the other decays, making it possible to ignore those dynamics, and focus solely on the exponential clearance of the latent cells.

#### Deriving the basic reproductive number *R*_0_ with the next-generation matrix method

Following Diekmann, Heesterbeek, and Roberts [50], we calculate the basic reproduction number of our complete model we begin by breaking apart our model into 2 matrices in the ‘next-generation’ (NG) fashion. We assume that the model is at the uninfected steady state so that 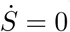 and define *S* = *S*_0_ = *α*_*S*_ */δ*_*S*_. Then, in matrix notation, calling **x** = (*L, A, V*)^*T*^ we write

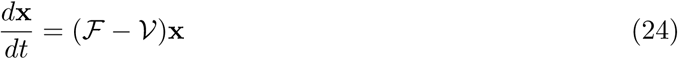

where *ℱ* is the matrix that describes new infections in each compartment

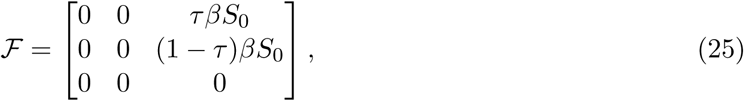

and *𝒱* is the matrix that describes the rates for leaving each compartment and for moving between them

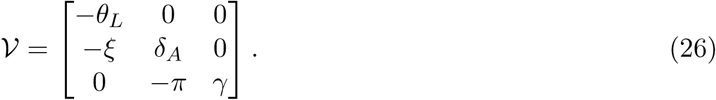

Then we want write the relationship

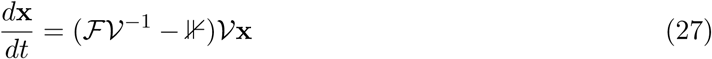

so that the largest eigenvalue of *ℳ*_NG_ = *ℱ𝒱*^−1^ will be our reproductive number, and we can see that at least 1 eigenvalue must be greater than 1 for an infection to take off. We have

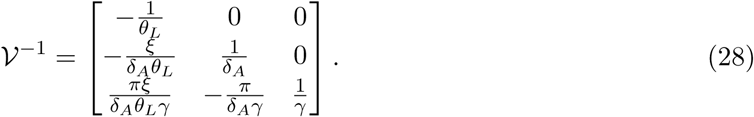

and thus

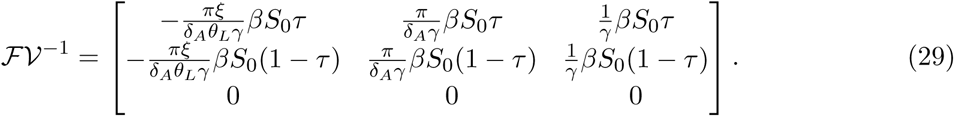

The next generation matrix admits two eigenvalues of zero and one that is positive. The positive eigenvalue is the largest eigenvalue of *ℳ*_*NG*_ and therefore we have the basic reproductive number of our model:

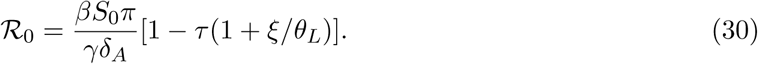

Comparing this value to the approximate derivation above (Eq. 17), we see that because ξ/*θ*_*L*_ *<* 1, the approximation is fairly accurate. Additionally, we could have carried through the calculation using infectivity as altered by ART efficacy without loss of generality.

#### Allowing direct active to latent transitions has a neglible impact on reservoir size

The contribution of actively infected cells to latency can be ignored such that we can consider the latent cell equation alone. This follows from a heuristic argument made by Conway and Perelson [29]. On ART, virus can be created by latent cell activation, which occurs rarely. However, the infectivity of HIV is reduced by *ϵ*_*c*_ on ART such that most susceptible cells are protected from infection. Further, the likelihood of any new infected cell becoming latent is quite small (*τ*). Thus, the product of the rare events–activation, new infection, and latency–is effectively zero.

Still, in some modeling works a transition from active to latent cells has been included [44, 52]. If we add a transition term to the model as a rate *φ*, the equations for the latent and active pool (decoupled from the virus and susceptible as before) are

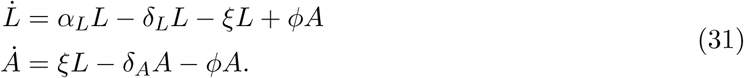

We verified numerically that only for unnaturally large values of *φ > α*_*L*_ (or likewise *φ > δ*_*L*_) does including this transition impact the reservoir size. In that case, there is a transient increase in the initial number of latent cells before the typical latent clearance dynamics take hold. This result is a consequence of the rapid death rate of active cells relative to the other rates.

#### Impact of duration and rates on clearance time

In the main article, we proposed that optimal latent clearance is achieved by increasing the duration of therapy rather than increasing the potency of the therapy equivalently (Fig. 2). If we examine the remaining fraction of latent cells over time *L/L*_0_ = exp [(*α*_*L*_ − *δ*_*L*_ − *ξ*)*t*], we can study the effect of multiplying any rate by a factor r, denoted *L*^(*r*)^ or multiplying the duration of time by factor *d*, denoted *L*^(*d*)^. The ratio of the percent remaining after doing either of these multiplications tells us which procedure is more valuable. For example, if we multiply the activation rate only,

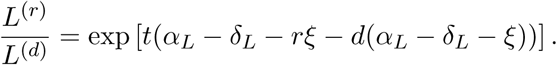

Then for *L*^(*r*)^ to be smaller than *L*^(*d*)^, we must have

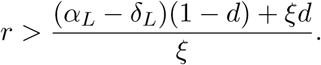

Note if we set *d* = r, we find that *d* < 1 because (*α*_*L*_ − *δ*_*L*_) is a negative number. Therefore, the decrease in the latent pool due to multiplying the duration of therapy will always be larger than an equivalent multiplication of the rate (unless the *r* < 1, which is a nonsensical proposition equivalent to making therapy less effective).

### Appendix II. Model parameters

#### Latent cell parameters: *θ*_*L*_, *α*_*L*_, *δ*_*L*_, *ξ*

Our most critical parameter values are those that describe the proliferation, death, and activation rates of latently infected cells–the sum of which is their net clearance rate *θ*_*L*_. Measuring these parameters *in vivo* in humans presents experimental challenges, and results depend on the models chosen to interpret the experimental data [53]. We acknowledge that there is uncertainty in these parameter estimates and thus provide an uncertainty and sensitivity analysis to demonstrate theoretical ranges for plausible outcomes (Fig. 4). Note that despite the variability of proliferation estimates, for even the slowest proliferation estimates we have encountered in the literature, proliferation rates are still two orders of magnitude larger than the estimated activation rate.

We use the proliferation rate *α*_*L*_ from Macallan *et al.* who measured *in vivo* turnover with deuterated glucose labeling in memory T cells in healthy adults and found that on average, 1.5% of CD45R0^+^CCR7^+^ T cells (central memory) proliferate daily, whereas 4.7% of CD45R0^+^CCR7^-^T cells (effector memory) and 0.2% of CD45R0^-^CCR7^+^ T cells (naïve) proliferate per day, corresponding with exponential growth rates of *α*_*i*_ = 0.015, 0.047, and 0.002, respectively [54]. We use Macallan *et al.*’s estimates from “healthy” adults (*i.e.* HIV-negative) because this study includes proliferation estimates for the CD4^+^ T cell subsets of interest.

In fact, several experimental works have demonstrated similar rates of memory CD4^+^ T cell proliferation among HIV-negative and HIV-infected on long-term ART [55, 25]. Whereas these studies examine the entire pool of CD4 T cells, rather than the smaller pool of latently infected cells, there is no empirical evidence to suggest that proliferation rates of latently infected cells would be *lower* than their uninfected counterparts. Indeed, Chomont *et al.* demonstrated that systemic proliferation events are tightly linked to the size and composition of the latent reservoir [12]. These studies show that CD4^+^ T cell proliferation rates are similar among HIV-uninfected and HIV-infected persons on long-term ART. However, net proliferation or proliferation of naïve versus memory cell rates are measured rather than the effector and central memory subsets that are now recognized as being important reservoir subsets, and thus we adopted the Macallan *et al.* rates.

Note that in our model, as in all T cell proliferation references we cite, proliferation rates reflect a combined impact of clonal expansion and homeostatic proliferation, as these are not yet distinguishable experimentally.

To estimate the activation rate of latent cells, Hill *et al.* used values found in Luo *et al.* from several structured treatment interruptions and found that on average, 57 CD4^+^ T cells per day transition from latency to the activated state [56, 21]. Assuming that the reservoir contains 1 million cells on average, the rate of activation of a single cell per day *ξ* = 5.7 *×* 10^-5^ [21]. Note that the activation rate *ξ* is several orders of magnitude smaller than the proliferation rate *α*_*L*_.

The net clearance rate of the latent reservoir is estimated from Siliciano *et al.*’s quantitative viral outgrowth assay as *θ*_*L*_ = -5.2 *×*10^-4^ per day [1]. This result was corroborated by Crooks *et al.* [26].

#### Susceptible and active cell parameters: *α*_*S*_, *δ*_*S*_, *α*_*A*_, *δ*_*A*_

In the complete model, the production of CD4^+^ T cells from the bone marrow and thymus is described by *α*_*S*_ ; and *δ*_*S*_ is the rate of susceptible T cell death. In several early HIV modeling papers, a value of 10 per *μ*L-day was estimated for the production rate [45, 15] with *δ*_*S*_ estimated at 0.02. Luo *et al.* used Bayesian statistical modeling to estimate HIV model parameters using data from 10 patients who underwent a series of 3-5 ART treatment interruptions with viral loads taken three times weekly following interruptions and then weekly following initiation of treatment [56]. They estimated *α*_*S*_ = 295, *δ*_*S*_ = 0.18, and *δ*_*A*_ = 1. Huang *et al.* also use Bayesian methods (Markov Chain Monte Carlo) and fit their model to data from Ref. [57], an AIDS clinical trial comparing dosing regimens for indinavir and ritonavir [58]. Huang *et al.*’s model also incorporated adherence, drug concentrations, and drug susceptibilities. They find *α*_*S*_ = 98.1, *δ*_*S*_ = 0.08, and *δ*_*A*_ = 0.37. Note in Table 2 that the ratio of *α*_*S*_ to *δ*_*S*_ varies between 500–1639 with the more recent experiments in relative agreement near 1500. Thus, we chose *α*_*S*_ = 300, *δ*_*S*_ = 0.2 to reflect this ratio.

**Table 2:** Proliferation and death parameters for susceptible and actively infected cells.

| Parameter | Perelson [45] | Huang [58] | Luo [56] | Markowitz [61] | Units |
| --- | --- | --- | --- | --- | --- |
| $\alpha_S$ | 10 | 98.1 | 295 | - | per $\mu\text{L}$-day |
| $\delta_S$ | 0.02 | 0.08 | 0.18 | - | per day |
| $\alpha_S/\delta_S$ | 500 | 1226 | 1639 | - | per day |
| $\delta_A$ | 0.24 | 0.37 | 1.0 | 1.0 | per day |

The death rate of productively infected cells *δ*_*A*_ was initially thought to be 0.24 [45]. Later, Perelson *et al.* find *δ*_*A*_ = 0.5 based on frequent sampling of 5 patients after giving ritonavir monotherapy [59]. Using a fixed viral clearance rate *γ* = 23 from [60], Markowitz *et al.* determined *δ*_*A*_ = 1 based on potent antiretroviral therapy with lopinavir-ritonavir, tenofovir, lamivudine, and efavirenz in 5 chronically-infected patients [61]. We chose *δ*_*A*_ = 1.0 to reflect more recent experimental findings. We assign *α*_*A*_ = 0 because the relative rate of proliferation of actively infected cells likely occurs at negligible rates compared to the death rate of these cells.

#### Estimating the infectivity *β*

Perelson *et al.* calculate the infectivity (using physical diffusion) of a virus finding *β* = 2.4*×*10^-5^*μ*L per day [45]. This value is ubiquitous but Luo *et al.* and Huang *et al.* also estimate parameter values for *β* as 3.9 × 10^-3^ and 1.7 × 10^-5^, respectively [56, 58]. Given that the range of these values spans two orders of magnitude, we chose a value between these extremes: 10^-4^.

#### Viral parameters: *π*, *γ*

The viral ‘burst rate’ *π* is the amount of virus a single actively infected cell emits in a day (roughly its lifetime). Rong and Perelson note that *π* is problematic, as its experimental value is under question and affects the values of many of the other parameters [44]. Haase *et al.* used radioactively-labeled RNA probes and quantitative image analysis to determine the number of viral particles per mononuclear cell in biopsies from fixed lymph tissue finding a mean value of 74 [62]. Using quantitative, competitive, real-time PCR to measure the mean viral RNA copy number per infected cell from fresh-frozen cervical lymph nodes from 9 HIV patients with varied viral loads, Hockett *et al.* found *π* = 10^3.6^ = 4 *×* 10^3^ [63]. Both of these estimates do not necessarily reflect the number of copies produced in a day per cell or in a cell’s lifetime, rather the amount that was being produced at the instant the experiments were performed. We adopt *π* = 10^3^. For the viral clearance rate *γ*, we use Ramratnam *et al.*’s estimate *γ* = 23, obtained from viral load measurements taken over 5 days before, during, and after apheresis in 4 patients assuming a constant rate of viral production [60].

#### The latency fraction: *τ*

The ‘latent cell fraction’ *τ* is the rate at which newly infected cells join the latent cell pool, whereas 1 − *τ* is the rate at which they join the actively infected pool. The few estimates of this parameter fall throughout a wide range. Despite this, the choice of *τ* within the given range does not affect our cure estimates or our estimate of the critical epsilon (*ϵ*_*c*_). Because Conway and Perelson’s estimates are based on modern measurements of the reservoir, we chose *τ* = 10^-4^, the upper bound on their estimates [29].

#### Parameter estimates, ranges, and confidence intervals

See Table 6 for the complete table of parameters that we incorporated in the uncertainty analysis.

**Table 6:** A summary of all parameters used in our simulations. \**δ*_*L*_ is back-calculated from known *α*_*L*_, *θ*_*L*_, and *ξ*. 95% confidence intervals are given in parentheses () where applicable from experimental data. Otherwise, the range is taken from our literature search or from the ranges given in the cited works not assumed to be normally distributed; these values are given in square brackets [].

| Param. | Value | (CI)/[Literature Range] | Dimensions | Source |
| --- | --- | --- | --- | --- |
| $\alpha_S$ | 300 | [10, 760] | cells per $\mu\text{L}$-day | [56, 45, 58] |
| $\delta_S$ | 0.2 | [0.02, 0.45] | per day | [56, 45, 58] |
| $\alpha_{cm}$ | 0.015 | (0.01, 0.02) | per day | [54] |
| $\delta_{cm}$ | 0.0155* | | per day | calculated* |
| $\alpha_{em}$ | 0.047 | (0.038, 0.057) | per day | [54] |
| $\delta_{em}$ | 0.0475* | | per day | calculated* |
| $\alpha_n$ | 0.002 | (0.0015, 0.0023) | per day | [54] |
| $\delta_n$ | 0.0025* | | per day | calculated* |
| $\delta_A$ | 1.0 | (0.8, 1.2) | per day | [56, 61] |
| $\xi$ | $5.7 \times 10^{-5}$ | $(5.4, 6.0) \times 10^{-5}$ | per day | [21] |
| $\theta_L$ | $-5.2 \times 10^{-4}$ | $-(2, 8.4) \times 10^{-4}$ | per day | [1] |
| $\beta$ | $1 \times 10^{-4}$ | $[0.016, 6] \times 10^{-3}$ | $\mu\text{L}/\text{copy-day}$ | [56, 45, 58] |
| $\tau$ | $10^{-4}$ | $[10^{-6}, 10^{-3}]$ | unitless | [29, 64, 15] |
| $\pi$ | $10^3$ | [20, 5340] | copies/cell-day | [56, 45, 58] |
| $\gamma$ | 23 | [9, 36] | per day | [60] |
| $L_0$ | $10^6$ | $[10^4, 10^8]$ | cells | [1] |
| $L_{cm}(0)$ | $0.6L_0$ | $[0.2, 0.8]L_0$ | cells | [12, 23] |
| $L_{em}(0)$ | $0.4L_0$ | $[0.2, 0.8]L_0$ | cells | [12, 23] |
| $L_n(0)$ | $0.02L_0$ | $[0, 0.1]L_0$ | cells | [12, 23] |

### Appendix III. Mycophenolate mofetil (MMF)

#### Determining the Hill coefficient and IC50s

In dose-response relationships, the concentration of drug that inhibits the response by 50% is called the IC50. The slope at the steepest point along the dose-response curve is called the Hill slope (or coefficient)[65].

The Hill coefficient and the IC50s for MMF were calcuated using the drc package in the R statistical computing language. Specifically, the 1 `drm` fitting command was used to fit the experimental *in vitro* proliferation data to the ‘LL.4’ (four parameter log-logistic function) function:

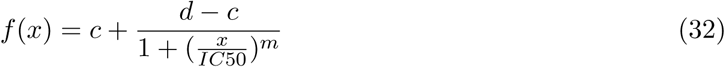

in which *f* (*x*) represents the fraction of proliferating cells at mycophenolic acid (MPA) concentration *x*. The upper limit of cellular proliferation in the experiment is *c* (no drug) and *d* is the lower limit (maximal drug effect)[47].

The active metabolite of MMF is mycophenolic acid, which was used in the titration experiments.

#### Previous studies in HIV-infected patients treated with ART and MMF

MMF has been given to HIV-infected patients in various settings, either experimentally as an antiviral drug or as part of standard regimens after kidney transplantation. Below we review the studies in which MMF was given to HIV-infected patients and either viral load reduction, reservoir reduction as measured by viral co-culture (QVOA), or time to viral rebound after treatment cessation were measured. Two trials assessed reservoir reduction using viral co-culture. Chapuis *et al.* revealed a reduction of the viral reservoir in the ART and MMF combination group but no reduction in the group on ART alone [25]. Sankatsing *et al.* also studied MMF and ART combination treatment and found a mean daily decay rate of latently infected cells (using viral co-culture) of 0.017 infected cells/10^6^ cells in patients on MMF and 0.004 infected cells/10^6^ cells in patients on ART alone. Despite the fact that this trend was not statistically significant(only eight patients in the MMF + ART group and nine in the ART-only group), the reservoir decay rate in the MMF-treated patients was almost five times as high as in the ART-only patients.

Both García *et al.* and Millán *et al.* demonstrated a delay in viral rebound after cessation of ART in patients who received MMF in addition to ART of 2-3 weeks [27, 39].

Margolis *et al.* gave MMF to five HIV-infected patients failing antiretroviral therapy with an average viral load of 10^4.78^ copies/mL. Four of the five patients experienced a greater than 0.5 log decrease in viral load compared to entry whereas three of five sustained the 0.5 log decrease after one year on ART combined with MMF.

Jurriaans *et al.* published a case of an HIV-infected patient who received a five drug ART regimen plus MMF and sero-reverted [40].

## Acknowledgements

We thank Keith Jerome for his helpful reading of the manuscript, the VIDD faculty initiative at the Fred Hutchinson Cancer Research Center, and the NIH for grants R01 AI116292 to FH, 1DP2DE023321-01 to MP, and U19 AI096111 and UM1 AI12662 to JTS. We also thank Claire N. Levy and Fernanda Calienes for assisting in the mycophenolic acid experiments. The following reagent was obtained through the NIH AIDS Reagent Program, Division of AIDS, NIAID, NIH: CEM CD4^+^ T cells from Dr. J.P. Jacobs.

### Author contributions statement

FH and JTS posed the initial question to model the effect of anti-proliferation on HIV latency; DBR, ERD, and JTS developed the computational model; SMH and FH devised, and SMH performed, the *in vitro* mycophenolic acid experiments; DBR and ERD performed calculations and produced figures; ERD, MP, FH, JTS performed literature review for parameter values. DBR, ERD, MP, FH, SMH, and JTS wrote the manuscript. DBR and ERD contributed equally to this work. FH and JTS also contributed equally to the work.

### Additional information

#### Competing financial interests

The authors declare no competing financial interests.

